# Attractor dynamics underlie sequence working memory in data-constrained recurrent neural networks

**DOI:** 10.64898/2026.09.30.755565

**Authors:** Matthijs Pals, Jakob H. Macke

## Abstract

Sequences of stimuli can be maintained in working memory by encoding each item, together with its position, as a distinct pattern of neural population activity. Competing hypotheses about the underlying dynamics have been proposed: patterns of activity might be supported either by stable attractor states that persist over time, or by transient sequential dynamics. To investigate the dynamics underlying sequence memory, we developed a framework for fitting low-rank recurrent neural networks (RNNs) to multi-session recordings of spiking activity, and applied it to 2,488 units recorded across 30 sessions in two macaques. Our data-constrained RNNs generated population spiking activity which closely resembled neural data and recovered single-unit tuning properties across sessions. Consistent with prior work, fitted networks represented items presented at different sequence positions in separate low-dimensional subspaces in the neural activity, alongside a dominant temporal component. The RNNs enabled direct investigation of the underlying dynamics: By isolating the temporal and stimulus subspaces, we found a manifold of fixed points emerged during the memory retention period. Targeted perturbation analyses confirmed that activity within this manifold is consistent with attractor-like dynamics, and predict that brief optogenetic perturbations delivered along this attractive manifold can selectively induce persistent shifts in neural activity and decoded behavior. Together, our results indicate that sequence working memory is organized by position-specific coding subspaces, within which attractor states maintain persistent representations, while activity evolves along time-varying dimensions.

## Introduction

Many behaviors require remembering ordered sequences, for instance holding a multi-digit authentication code in mind long enough to enter it correctly. Working memory is the capability that allows holding and manipulate information over short periods during such tasks [1]. Early theories proposed that working memory was instantiated by the persistent firing of neurons throughout the memory interval [2–4]. In addition to reports of persistent activity (e.g., [5, 6]), subsequent work has revealed that many individual neurons exhibit complex, time-varying responses [7–11]. One way to reconcile these observations is that the information can remain stable at the population level during memory maintenance, [12–14].

In sequence working-memory tasks, population activity exhibits additional structure: Items presented at different serial positions can occupy distinct low-dimensional subspaces within the population activity, as observed in experiments with both single unit resolution [15–17] and broadband MEG recordings [18]. This geometry resembles a representation in which items can be bound to sequence position along (approximately) orthogonal axes [19, 20]. Such a scheme enables ordered information to be maintained within a single population despite overlapping neural units.

However, representational geometry alone does not reveal the dynamical mechanisms that underlie these patterns. Multiple competing hypotheses remain under active investigation in the field [21]; the same stable population-level representation could arise either from persistent attractor states [14, 22–28] or from sequences of decaying transients [29–31]. We therefore ask: what dynamical regime underlies memory maintenance in the observed sub-spaces during working memory for sequences?

To address this question, we design a framework for fitting recurrent neural networks (RNNs) at single trial resolution to multi-session single-unit recordings, which we apply to data of macaques performing a sequence working-memory task [16]. RNNs are widely used to model neural population activity and to generate hypotheses about the computations they implement [32–37]. Here we focus on RNNs with low-rank connectivity [38–43], which provides interpretability through a direct link between high-dimensional population activity and an underlying low-dimensional dynamics

Our fit low-rank RNNs are generative models, that capture both spiking statistics and recover experimentally observed population level representations of 2,488 units recorded across 30 sessions in two macaques. We perform stability analyses of the low-rank RNNs latent dynamics and in silico perturbation experiments. These analyses show that the network maintains sequence information via stable attractor dynamics together with a time-varying component. Our results predict that transient perturbations delivered along the memory manifold during the delay period will induce persistent shifts in the neural state, resulting in behavioral changes.

## Results

### Dynamical mechanisms underlying stable working memory representations

To study the dynamics underlying working memory, we use a dataset of single-unit recordings from the macaque prefrontal cortex (PFC) [16]. Macaques maintain sequences of 1 to 3 stimuli which, after a delay of 1.15–1.5s, are indicated by pressing on the screen (Fig. 1a). Chen et al. [16] found that each item in the sequence was represented in approximately orthogonal subspaces of the population activity (schematically in Fig. 1b), in line with other recent studies [15, 17, 18]. These subspaces can be consistently decoded throughout the memory period (schematically in Fig. 1c).

**Figure 1.**
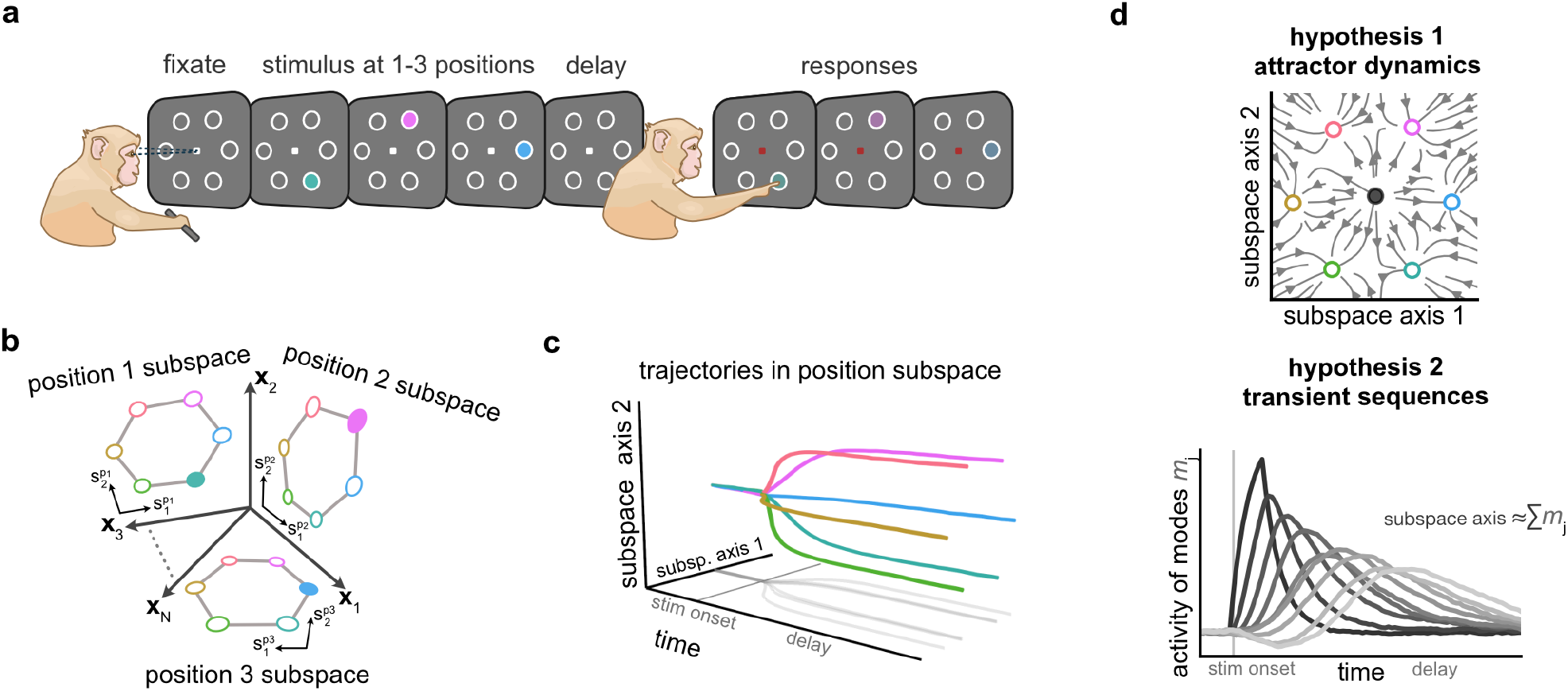
Two hypotheses underlying population codes during working memory. **a)** We use data from an experiment in which macaques performed a task requiring them to maintain sequences of 1–3 stimuli in working memory [16]. **b)** Previous studies found a separate (approximately orthogonal) subspace for each sequence position in the population activity of PFC neurons [15–17]. **c)** Subspaces are consistent throughout the delay. **d)** Stable decoding during the delay period could arise from either attractor dynamics (top; [14, 27, 28, 41]) or transient sequential activity (bottom; [29–31]).

Stable decoding of remembered items during the delay could arise from distinct dynamical mechanisms (Fig. 1d): one possibility is that *attractor dynamics* underlie persistent activity patterns, for instance via a fixed point for each stimulus (Fig. 1d, top) [14, 22, 27, 28, 41]. Alternatively, the observed time-invariant subspace could emerge from *transient sequential activity*, in which a sequence of modes (patterns of activity) that trigger one another bridge the delay (Fig. 1d, bottom) [29–31]. Summing these transient modes can also result in a stable constant representations throughout a memory period [29]. To formalize these hypotheses, we constructed RNN models that maintain a stable memory representation while using either attractor or transient dynamics (Fig. S.1, S.2). While we observed a similar geometry of the latents between both models, we confirmed that our subsequent analyses can distinguish between the two models (Fig. S.3, S.4).

### Data simulated with multi-session RNNs match key properties of recordings

To distinguish between the mechanisms proposed above, we train generative RNN models of the spiking activity of macaques. Because the recordings are collected over multiple days, we employ multi-session low-rank RNNs that capture shared population dynamics (Fig. 2a). Each recorded unit is mapped to a corresponding RNN unit (Fig. 2a, left), and the low-rank constraint on recurrent weights (Fig. 2a, middle) ensures that network activity evolves within a shared, low-dimensional subspace that is shared across sessions (Fig. 2a, right). We use RNNs with piecewise-linear (ReLU) activation functions, allowing fixed points and other dynamical features to be efficiently identified [35, 44–48].

**Figure 2.**
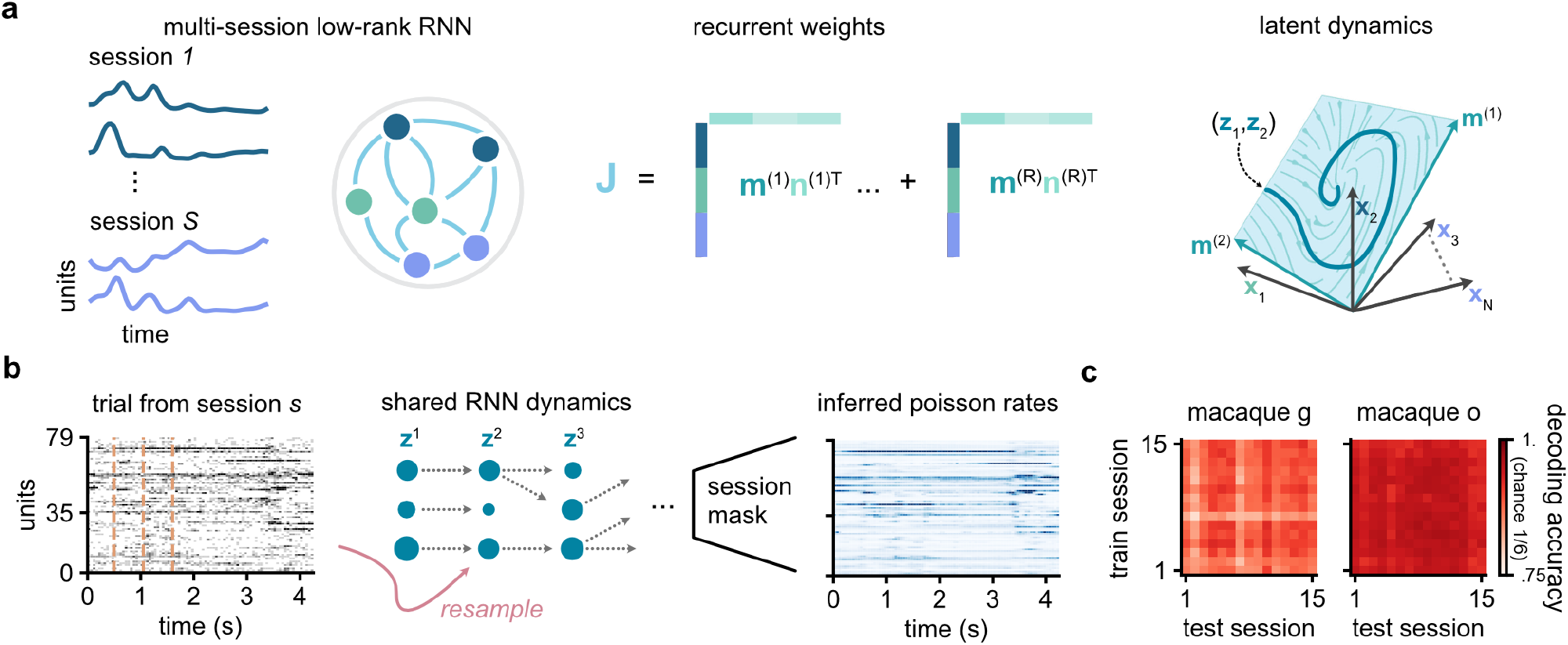
Multi-session low-rank RNN with consistent latents between sessions. **a)** Each unit in the RNN is identified with one recorded unit (left). We assume that the recurrent weights have a low-rank structure (middle), meaning that we can write the RNN’s recurrent connectivity weights **J** as an outer product of left and right connectivity vectors **m** and **n**’s. The low-rank connectivity structure implies that the multi-session neural activity is constrained to a shared low-dimensional subspace (right; spanned by the **m** and input vectors). **b)** To fit the RNNs to neural data, we use sequential Monte Carlo methods, and parameterized latent dynamics by a low-rank RNN that is shared between sessions. During training we alternate trials from different sessions. **c)** The latents between sessions are consistent with each other as shown by using the inferred latents during the late delay period to do cross-session decoding of the maintained items.

To fit these RNN models at single-trial resolution, we develop a method for fitting low-rank RNNs to multi-session data (Fig. 2b). This method builds on a previous framework Pals et al. [48] and leverages sequential Monte Carlo methods [49–52]. During training we alternate data from different sessions. We use a so-called bootstrap proposal: At each time step, we propagate multiple latent trajectories (‘particles’) forward according to the low-rank RNN dynamics. The sampled trajectories are then resampled based on how well they explain the observations in the currently presented session. Using this approach, we fit 5 rank-64 RNNs to 15 sessions of recordings of macaque G (1121 units, 10892 trials used for training) and 5 models to 15 sessions of macaque O (1367 units, 6852 trials). This rank was required to capture stimulus-conditioned delay responses (Fig S.5). To verify that latent dynamics are consistent between sessions after fitting the model, we perform a cross-session decoding analysis. We first use our inference setup to obtain posterior latents for each session separately, then train session-wise decoders to decode presented stimuli from late-delay latents. By testing each decoder on the inferred latents of all other sessions, we confirmed our framework results in a low-rank RNN that successfully captures a shared latent structure underlying data collected over multiple sessions (cross-decoding accuracies: macaque G: 0.87 *±* 0.009, 0.85 *±* 0.01, 0.90 *±* 0.01, macaque O: 0.97 *±* 0.01, 0.94 *±* 0.01, 0.92 *±* 0.009, mean *±* sd over 5 models, for sequence position 1 to 3, respectively; Fig. 2c).

After fitting the RNNs, we used them to generate new trajectories of latent dynamics and corresponding spiking activity for all units in the data, given stimulus inputs (Fig. 3a). The generated spiking activity shows close correspondance with the recorded activity across held-out spiking data from all sessions, both in terms of firing rates (macaque G: *r* = 0.996 *±* 0.008, macaque O: *r* = 0.999 *±* 0.001), inter-spike-interval (ISI) variability (G: *r* = 0.980 *±* 0.01, O: 0.984 *±* 0.006), and pairwise correlations between units (G: *r* = 0.888 *±* 0.04, O: *r* = 0.895 *±* 0.02; all *µ ±* sd over 15 sessions, after first averaging the *r* values over 5 trained RNNs per macaque; Fig. 3b). Previous reports [15] have found units that jointly code for position and stimulus, i.e., fire in response to a stimulus only when it is shown at a particular sequence position, and might be responsive to different stimuli at different sequence positions. Such units are also found in our model (Fig. 3c; see also Fig S.5 for additional statistics).

**Figure 3.**
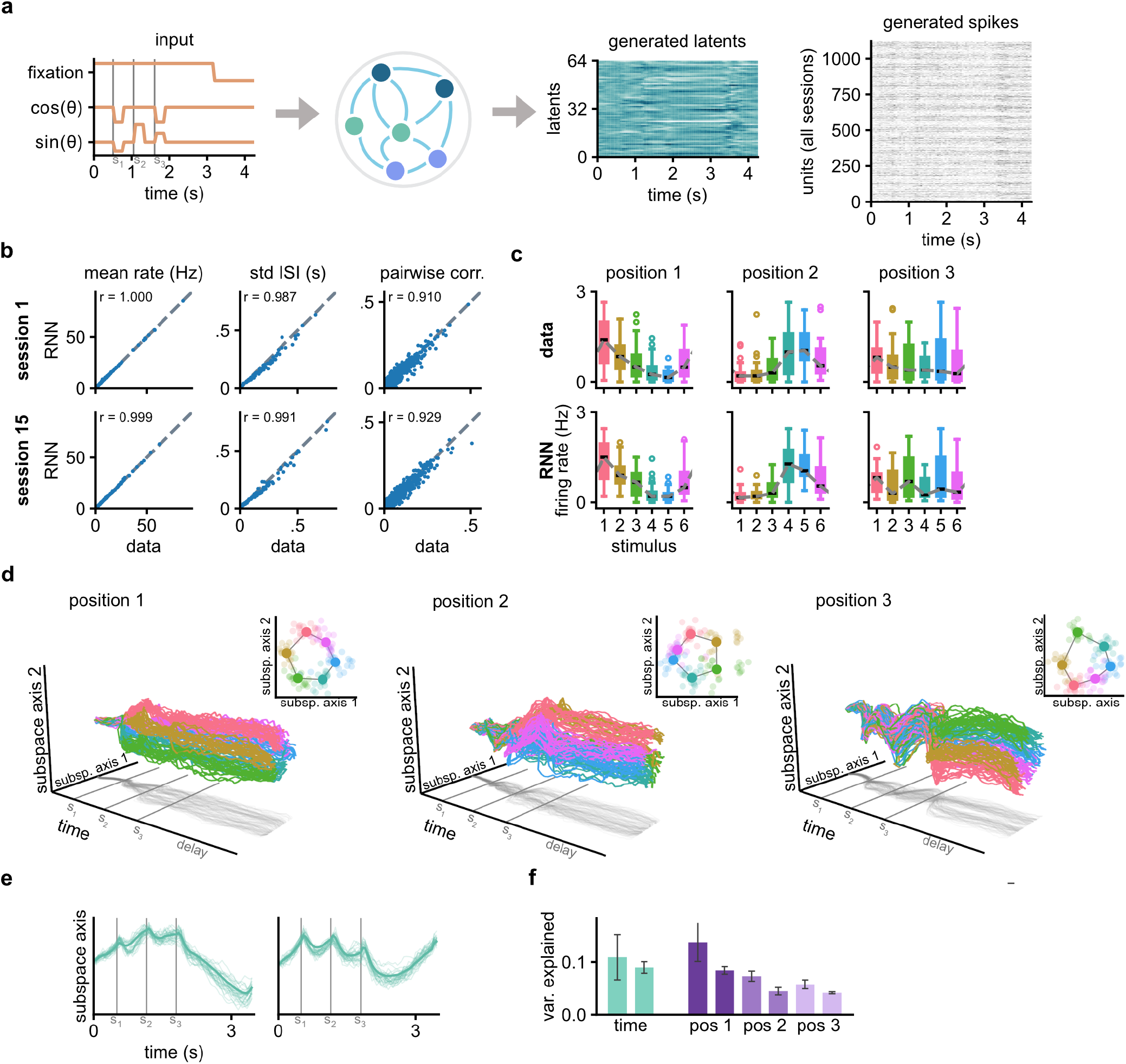
Generative RNNs capture key properties of the data. **a)** Our data-fitted RNNs can be used as generative models that produce stochastic spike trains for all recorded units, conditioned on task inputs: A constant fixation input until the end of the delay, and the sequence of task stimuli to be maintained (encoded as sin(*θ*), cos(*θ*), where *θ* is the angle of the stimulus). **b)** Key statistical properties of neural activity are accurately reproduced by the model. Comparisons between RNN-generated data and held-out recordings show agreement in mean firing rates, inter-spike-interval (ISI) variability, and pairwise correlations between neurons. Two example sessions are shown. **c)** Units with mixed selectivity are captured by our RNN as shown by this example tuning curve of a neuron that jointly encodes stimulus identity and sequence position. **d)** Orthogonal transformations of the latent activity reveal three stimulus-encoding subspaces, each corresponding to one sequence position. Neural trajectories within each subspace separate according to the stimulus presented at that position (colors); each line represents the activity in one trial. **e)** Additional latent dimensions capture temporal dynamics that are largely independent of stimulus identity, including ramping activity during the delay and transient responses at stimulus onset. **f)** Identified latent dimensions consistently explain a large amount of delay activity across model seeds and macaques.

For population level analysis, we first find an orthonormal transformation of the RNN dynamics that allows us to find directions in RNN activity space that capture task-related variability [14]. Concretely, we find orthogonal two-dimensional subspaces for each sequence position: one in which the first item in the sequence is encoded, one for the second, and one for the third. This organization is in line with previous experiments [15, 16] and models [20, 53]. Additionally, we also find components in the latent dynamics that correlate with timing information [7, 54], such as stimulus onset and progression of the delay period (Fig. 3e; hereafter referred to as ‘time-varying latents’). To quantify to what degree the neural activity during the delay can be explained by stimulus and timing information, we compute the variance explained by each of the found basis vectors (Fig. 3f). In total the 8 basis directions explain 64% *±* 1.5% (*±sd n* = 10) of variance of the RNN’s delay activity, indicating we have isolated relevant directions of neural activity during the memory period.

### Analysis of RNNs indicates attractor-guided dynamics

Having verified that our model generates activity that matches the recordings, we next analyse the RNNs dynamics. The time-varying activity observed during the delay period implies that the network state does not converge to a fixed-point attractor during memory maintenance. Yet, time-varying activity does not rule out attractor dynamics [55], for example, chaotic or periodic attractors have been suggested to underlie neural representations during working memory [56, 57]. Alternatively, previous work suggests that activity may evolve in directions orthogonal to the subspace in which stimulus information is maintained by static attractors [13, 14].

In particular, the independence between the memory subspaces and the time-varying component described by Murray et al. [14] suggest that the dynamics within the memory subspaces can be studied separately. To do so here, we simulate the RNN until it is near the end of the delay period. Then, we clamp the time-varying latent variables to their mean value (across all conditions), and continue the simulation for several additional seconds (Fig. 4a), while keeping the fixation input on. In this ‘clamped’ system, we find that models tend to converge to a steady state. To identify the corresponding fixed points of the clamped system, we apply the algorithm of Eisenmann et al. [46]. We find both stable and saddle fixed points that cluster around the mean latent values during the delay period (Fig. 4b), for models trained on data of both macaques, irrespective of the exact clamping timepoint (Fig. 4c). This result suggests that the dynamics during the delay may be guided by an underlying attractor structure.

**Figure 4.**
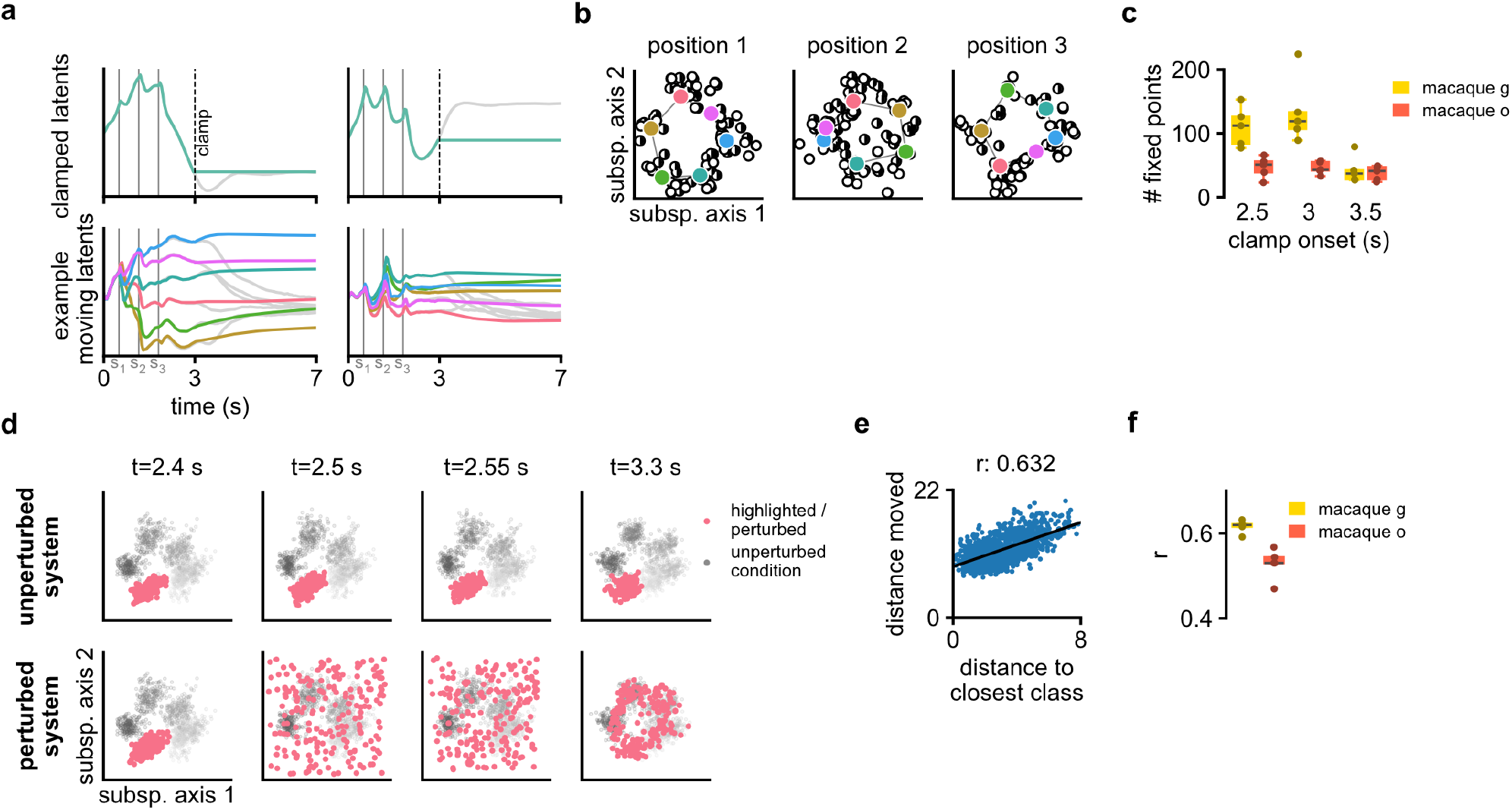
Signatures of attractor dynamics. **a)** To isolate attractor dynamics in the presence of time-varying activity, we clamped the time latents to their mean value near the end of the delay period and simulated the remaining latents. Grey trajectories show how latents evolve when not clamped. **b)** Analysis of the clamped latent system reveals multiple stable fixed points (white) and saddle points (half-black/half-white) that cluster around the average neural activity observed during the delay periods (colored circles), suggesting that trajectories are guided toward a stable manifold in this region of state space during memory-maintenance. **c)** We find multiple fixed points for all models, for a range of different timepoints used for clamping. **d)** Activity in one memory subspace during the delay, unperturbed (top row) and with a perturbation applied at the start of the delay (bottom row) in all trials of a highlighted condition (pink). Perturbations converge back to the hexagonal structure, indicating this is a stable manifold in this subspace. **e)** Quantification of perturbations (like in c, but for all conditions). In line with an attractive manifold, perturbations that land closer to one of the condition means (on-manifold) show less movement than those that land further away (off-manifold), which we can quantify by an *r* value. **f)** We compute this *r* value for all models fit to either macaque. We can consistently find indications of attractor dynamics across seeds for both macaques.

Different dynamical regimes can also be distinguished by their responses to perturbations ([22, 58, 59]; responses to perturbations in hand-constructed models with either attractor or transient dynamics in Fig. S.1, S.2). We apply perturbations to the RNN activity within one of the stimulus subspaces, at the start of the delay period, and examine how the activity evolves thereafter. If the ring of activity reflects a manifold of stable states, perturbations that remain on the manifold should produce little subsequent movement, whereas perturbations that move the activity away from the manifold should decay back toward it, and therefore exhibit larger movement. In contrast, if the subspace were marginally stable during the delay, perturbations in any direction should produce little movement. Finally, if the subspace corresponded only to an output dimension rather than the internal subspace where stimuli are stored, all trajectories should decay back toward the original state. The behavior observed in (Fig. 4d-f) is most consistent with the presence of a stable attractor manifold, as the perturbed trajectories relax back to the ring structures in the memory subspaces.

Finally, we use the model to generate experimental predictions: We ask if we can use targeted perturbations during the memory period to change the macaques’ response for one sequence position, while keeping its response to the other positions unaffected. To answer this question, we first identify a neural subspace from which we could decode behavioral responses, using our RNNs to obtain posterior latents (with sequential Monte Carlo) matched to macaque behavioral outcomes. The identified subspace can subsequently be used to obtain behavioral responses from activity generated by simulating the RNN; We can decode the sequence of stimuli presented to the RNN with an accuracy of 0.87 *±* 0.029, 0.77 *±* 0.034, 0.72 *±* 0.051 for sequence position 1, 2 and 3 respectively (*µ ± sd, n* = 10). Next, we apply targeted perturbations to the previously identified stimulus subspace, which produce persistent shifts in the network state that lead to corresponding changes in the decoded behavioral output (Fig. 5). To test whether these stimulus subspaces can be causally manipulated using experimentally relevant interventions, we apply a perturbation that is meant to approximate optogenetic-style perturbations during the delay period. Specifically, we inject the same constant current into a subset of units associated with the target stimulus subspace (note that this assumes that neurons contributing to the relevant subspace can be selectively identified and targeted). Because these perturbations cannot be perfectly aligned with the low-dimensional position subspaces, their effects are somewhat less selective than idealized subspace-targeted perturbations, but nevertheless induce reliable shifts in the network state (Fig. 5b; right). These results suggest that a brief perturbation to a stimulus subspace during the delay can steer the macaque’s response toward a desired target outcome, specifically for one stimulus position.

**Figure 5.**
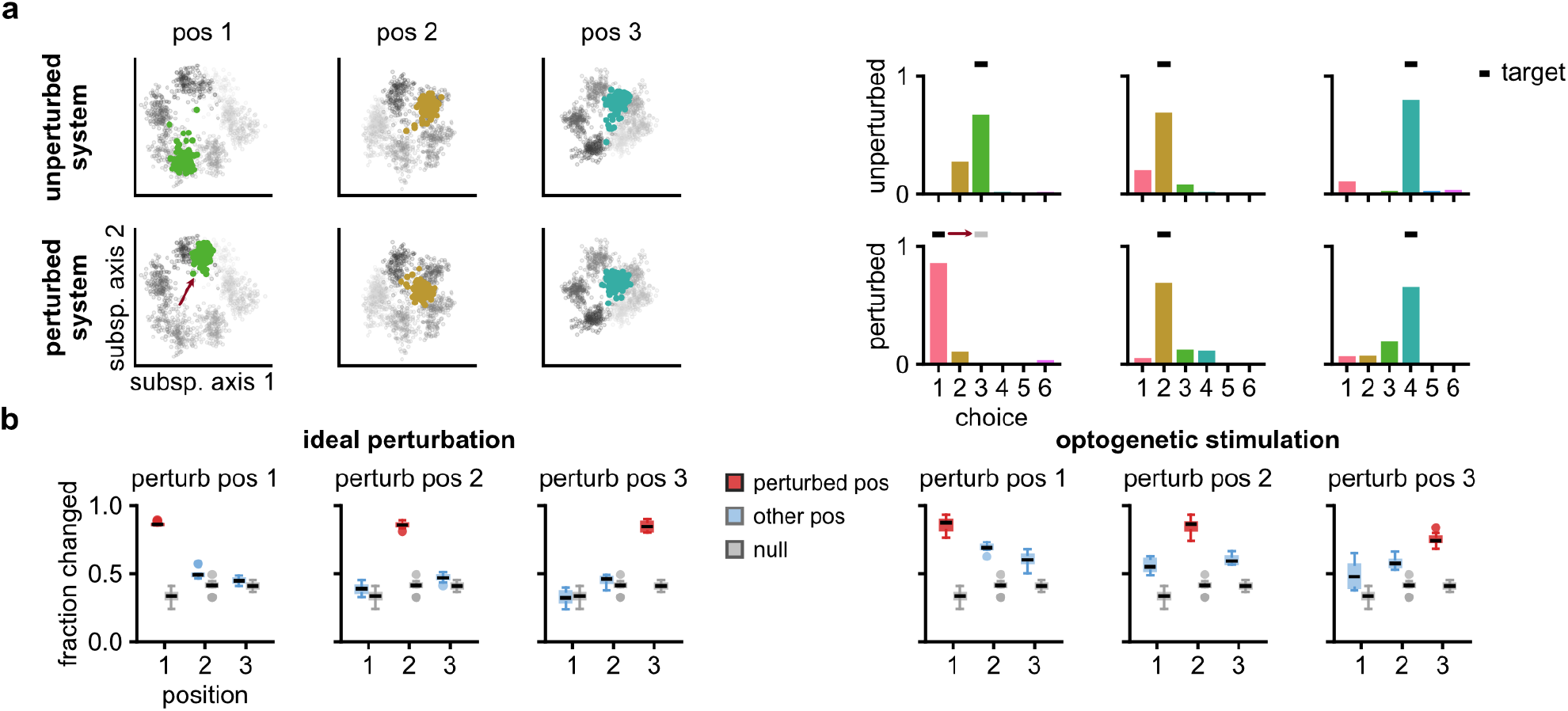
Targeted perturbations selectively change behavioral responses to one sequence position. **a)** To test whether the identified position subspaces could be relevant for behavior, we apply targeted perturbations that selectively change the models decoded responses for one particular sequence position. Here, we add a perturbation to the position 1 subspace (left), on multiple repetitions of the same trial (with sequence 3,2,4). We can change the model’s response from choosing item 3 to item 1 for the first sequence position, while leaving the responses for the latter two positions mostly unaffected (right). **b)** Across all possible permutations, and all trained models, we can consistently change the target position. We retain this effect if we use a perturbation where the same current is applied to a subset of units (mimicking optogenetic perturbations), although with a decreased position specificity.

## Discussion

Our results suggest that sequence working memory in macaque prefrontal cortex is supported by stable attractor dynamics embedded within a low-dimensional, time-varying population structure. We fit low-rank RNNs to multiple sessions of neural recordings of macaques performing sequence working memory tasks [16]. The obtained RNNs are generative models that can generate realistic spiking data with underlying latent trajectories matching the previously reported geometry [15–17]. Through both stability analysis and perturbation analysis, we find support for stable attractor dynamics next to time-varying delay activity.

We obtained these results using a new approach for fitting multi-session low-rank RNNs. This complements approaches such as LFADS [60], with a potential additional session-alignment loss (NoMAD; [61]). In LFADS, trial-to-trial variability is captured through initial conditions, per-trial biases [62], or optionally inferred inputs. In contrast, our approach learns stochastic transitions [48], facilitating a generative model of neural activity; related stochastic-latent approaches include XFADS [63]. Other recent multi-session approaches learn session-specific low-rank perturbations [64], hypernetwork-generated connectivity [65], or session-specific reweighting of a shared connectivity basis [66]. In contrast, our framework learns one dynamical system that jointly generates activity for all sessions.

Our findings connect two perspectives on working-memory dynamics that are often considered distinct: stable attractor dynamics and time-varying population activity. In our data-constrained models, stimulus information was maintained within stimulus-specific attractor manifolds, while activity evolved over time along orthogonal dimensions. This organization relates to dynamical architectures proposed in previous studies using task-trained or hand-designed models. Murray et al. [14] and Druckmann and Chklovskii [13] similarly proposed a separation between stimulus-related and time-varying dimensions, although stimulus representations in our models form stable, attractive circular manifolds, rather than the marginally stable planes considered in those studies. Likewise, Chaisangmongkon et al. [67] found stimulus-dependent attraction alongside time-varying activity in RNNs trained on working-memory tasks, with trajectories converging to what they termed ‘tunnels’. Our results extend these theoretical proposals by showing that a similar dynamical organization can account for population activity directly recorded from the brain.

An important question is whether the inferred attractor dynamics reflect the underlying system or biases of the model class. Recent work has highlighted potential identifiability issues when fitting RNNs to partially observed systems, showing that models can favor attractor-like solutions when approximating high-dimensional transient dynamics, particularly under limited capacity [68]. Here, we use comparatively large networks, which could mitigate this bias. More directly, we applied our fitting method to data generated from models with either transient or attractor dynamics. In these student-teacher setups, we can recover the correct underlying mechanism (Fig. S.3, S.4).

Transient and attractor dynamics make distinct predictions about how the system responds to perturbations or noise, as also noted by Qian et al. [68]. Perturbation approaches have recently been applied successfully in vivo to distinguish dynamical mechanisms [22, 59]. Our model makes clear predictions for how targeted perturbations should affect subsequent dynamics, and behavioral responses during sequence working memory tasks, providing a route for experimental validation. Analyzing trial-to-trial deviations from the mean dynamics [69] may offer an additional non-invasive way to distinguish between these regimes.

In summary, our results indicate that sequence working memory representations are supported by attractor dynamics within position specific coding subspaces, which coexist with the time-varying population activity observed during working memory. Future work combining data-constrained models with causal perturbations will be essential for verifying whether the attractor structure inferred is a property of the biological system.

## Acknowledgments

We are grateful to Prof. Liping Wang and Dr. Jingwen Chen for providing the macaque single unit recordings. This work was supported by the German Research Foundation (DFG) through Germany’s Excellence Strategy (EXC-Number 2064/1, PN 390727645) and SFB 1089 (PN 227953431), the European Union (ERC, DeepCoMechTome, 101089288) and the German Federal Ministry of Research, Technology and Space (BMFTR) under the NeuroAI initiative (grant no. 01GQ2509B, NAILIt). MP was a member of the International Max Planck Research School for Intelligent Systems (IMPRS-IS). We thank João Barbosa for feedback on the manuscript.

## Methods

### Code availability

The code for training RNN models, running our analyses and generating the paper figures is available at: https://github.com/mackelab/swm_rnn.

### Details of the data

We used data from Chen et al. [16], see the corresponding paper for in-depth details. In brief, macaques were set in front of a screen and performed a sequential working memory task. Each trial started with the macaque pulling a level and fixating at a yellow fixation square. After 100 ms 6 empty circles situated on the vertices of a hexagon appear, and the fixation square turned white. After another 500 ms one to three stimuli were presented sequentially for 250 ms each with an inter-stimulus interval of 300-500 ms. After a delay of 1150 to 1500 ms the white square turned blue, the macaques released the level and touched locations on the screen. If the touched locations matched the stimuli, the macaques received a drop of water or juice. Note that the experiment contained both recording days were the monkeys had to report the stimuli in the presented (forward) order, as well as days were they had to report them in reverse order. For this study we only use forward recordings.

Recordings were obtained using a semichronic microdrive system (Gray Matter Research, USA [70]) implanted in the left hemisphere. A 157-channel tungsten electrode array (LS-157, AlphaOmega; *∼* 1 MΩ, 1.5 mm interelectrode spacing) was used and the recording chamber covered the prefrontal and premotor areas. Electrode depths were adjusted to maximize the number of recorded units. High-frequency signals (300–8000 Hz) were sorted offline using IronClust and manually curated (Plexon Inc.). Both single- and multi-unit activity were included. Units were discarded if they had firing rates <1.2 Hz or on average less than 5 spikes per trial. We discretized the recordings to 50ms bins. We used the first 15 recordings from two macaques, giving us 1121 (macaque G) and 1367 (Macaque O) units.

### Recurrent neural networks

We use RNNs of the form (e.g., [33, 71, 72]):

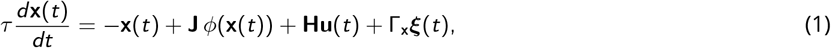

where **x**(*t*) ∈ ℝ^*N*^ is the vector of neural activity (currents), *τ* ∈ ℝ^+^ is the synaptic time constant, and **J** ∈ ℝ^*N×N*^ is the recurrent connectivity matrix. The task stimuli are represented by input 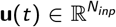 and corresponding weights 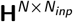, and ***ξ***(*t*) ∈ R^*R*^ is white noise input with coupling matrix Γ_**x**_ ∈ R^*N×R*^. The nonlinearity is applied element-wise as:

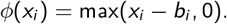

We assume that each RNN unit maps to one recorded unit, and that the recorded spiking activity can be described by an underlying continuous rate *r*_*i*_ (*t*). Since the RNN activity *ϕ*(**x**(*t*)) is dimensionless, we introduce a per-neuron gain and bias.

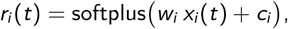

where the softplus(*x*) = log(1 + *e*^*x*^) ensures non-negative rates.

#### Low-rank structure

High-dimensional neural activity can be studied by looking at an underlying low-dimensional system [36]. This link can be made explicit in low-rank RNNs [40–43]. Low-rank RNNs also facilitate our multi-session inference setup, as we model units recorded over different sessions, whose activity then lives in a shared low-dimensional subspace. To construct these networks, we constrain the weight matrix **J** to have rank *R*:

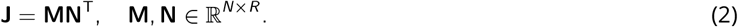

This results in the dynamics being confined to the linear subspace spanned by **M** and **H** (assuming **x**(0) in this space, or after initial transients have decayed, and Γ_**x**_ = **M**Γ_**z**_). This implies we can describe the RNN dynamics by an *R*-dimensional variable **z**(*t*) and an 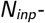 dimensional variable **v**(*t*); we can write

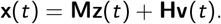

We can now equivalently write the RNN dynamics (Eq. 1) by describing how the activity evolves in the (*R* + *N*_*inp*_)-dimensional subspace:

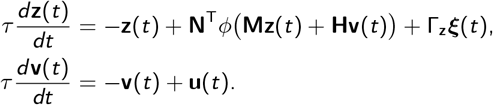

#### Discretization

We discretized the latent dynamics using the Euler-Maruyama method with timestep Δ_*t*_. Let 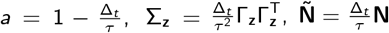, we obtain

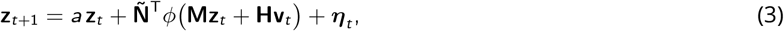

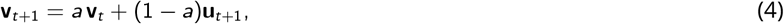

with ***η***_*t*_ *∼ N* (0, Σ_**z**_) and where, with a slight abuse of notation, we wrote **z**_*t*+1_, **v**_*t*+1_ for integrating the system one time-step ahead (i.e., for **z**_*t*+Δ*t*_ ,**v**_*t*+Δ*t*_). The currents of the RNN units, and corresponding predicted rates are given by:

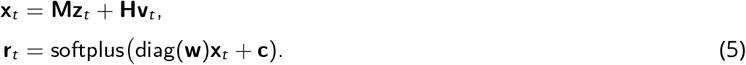

#### Probabilistic model

Given observed data **y**_1:*T*_, the generative model corresponding to:

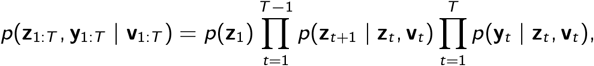

where **v**_1:*T*_ is deterministically defined from the input **u**_1:*T*_ by Eq. 4. The latent transition distribution is given by the low-rank RNN dynamic (Eq. 3):

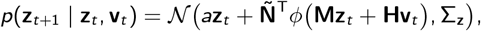

and we also assume the initial condition to be Gaussian distributed: *p*(**z**_1_) = *N* (***µ***_1_, Σ_1_). Observations are assumed to be Poisson distributed:

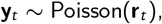

with the rate **r**_*t*_ given by Eq. 5.

### Inference and training

#### Sequential Monte Carlo

Because the exact posterior *p*(**z**_1:*T*_ | **y**_1:*T*_) is intractable, we approximated it using Sequential Monte Carlo (SMC) with a set of *K* particles representing latent trajectories [52]. At each timestep, latents were proposed from the bootstrap proposal, i.e., from the RNN dynamics *p*(**z**_*t*_ | **z**_*t*−1_) and assigned importance weights:

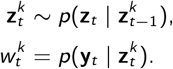

These were normalized as 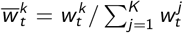 followed by a resampling step. This process results in a filtering approximation to the posterior at time *t*:

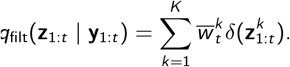

We optimized the RNN by maximizing an Evidence Lower Bound (ELBO) defined over the joint distribution of all particles and ancestor indices, *q*_smc_ [49–51]:

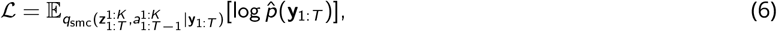

Where 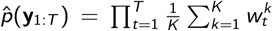 is the unbiased marginal likelihood estimate. During optimization we discarded high-variance gradient terms associated with the resampling step [49–51].

#### Optimisation

During training we alternated batches drawn from different recording sessions. For computing the loss, we only considered the rates given by RNN units matching the data units in the current batch. We minimised the ELBO (Eq.6) with stochastic gradient descent, using the Adam [73] optimiser in Pytorch [74].

We used batches of 128 trials, and trained for 750 epochs. Each training epoch consists of 15 sessions, with 344-571 (macaque O) and 566-850 (macaque G) trials per session. We used cosine decay to 10^−6^ for the learning rate, after an linear warm up of 50 epochs from 10^−6^ to 10^−3^. Training took around 8 hours on one NVDIA 3080 for macaque O and 12 hours for macaque G. There was a training instability on some runs; In case NaNs or divergent losses appeared, a run was restarted. The models had a rank of *R* = 64, and we used *k* = 32 particles.

#### Initialisation

Models were initialized as follows:

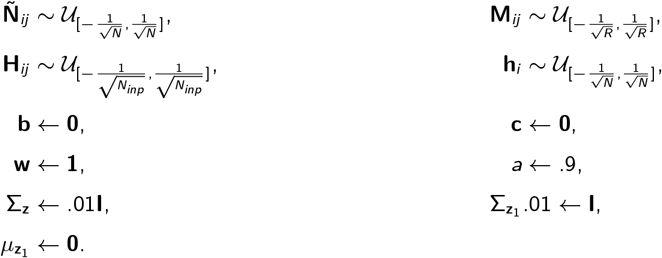

We constrained *a* to be between 0 and 1 by instead optimizing ã with the following (sigmoidal) parameterization *a* = exp(− exp(ã)) [75]. We assumed all covariances are diagonal and isotropic; parameterized by their standard deviations using 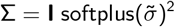.

### Stimuli and time basis directions

To obtain interpretable low-rank RNN dynamics, we expressed the latent activity in terms of orthogonal modes aligned with task variables. The connectivity vectors **M** were unconstrained during training and therefore do generally not define an orthogonal coordinate system. This means that individual latents **z**(*t*) can reflect arbitrary linear combinations of underlying modes. We obtained an orthogonal basis after training, by performing a singular value decomposition of the connectivity matrix,

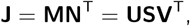

and projecting the latent state as **z** ← **U**^⊤^**Mz** (retaining only the first *R* singular vectors).

This basis is still agnostic to task structure. To better understand which latent dimensions encode variables such as stimulus identity we performed an additional orthonormal transformation that aligned components with task-relevant variables (inspired by Murray et al. [14]).

The latent activity was arranged as a tensor 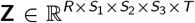, where *R* = 64 latents, *S*_*i*_ = 6 stimulus identities at each of three sequence positions, and *T* denotes time. Because not all stimulus combinations were valid (each item could appear only at one position) and sequence elements had unequal durations, we defined a binary mask *M* indicating valid samples.

We first subtracted the global mean activity of each neuron across all valid samples:

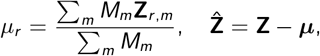

where *m* runs along all task and time dimensions.

To isolate task-related variance, we computed marginalizations for time and each sequence position. For a given marginal (e.g., position 1: *i* = *S*_1_), activity was averaged over all other dimensions using the mask:

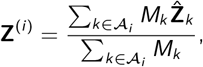

where *A*_*i*_ denotes all axes except neurons and the marginal of interest. Only samples with nonzero mask support were retained, and the result was reshaped to 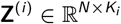.

We then performed PCA:

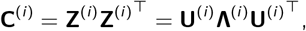

and the leading eigenvectors were retained to form **W**^(*i*)^. We kept 2 components for time and each of the three sequence positions.

Subspaces were extracted sequentially (time, position 1, position 2, position 3). After each step, the new basis **W**^(*i*)^ was orthogonalized with respect to previously extracted components *W*_prev_:

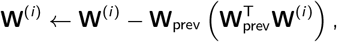

followed by normalization of the columns. To prevent variance leakage across marginals, we projected out the sub-space at the current step:

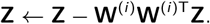

All **W**^(*i*)^ were concatenated to form **W**_enc_ ∈ R^*R×*8^. This basis was extended to a full orthonormal basis **W**_full_ ∈ R^*R×R*^ by appending random vectors and orthogonalizing the combined set via QR decomposition.

We computed the basis vectors **W**_full_ using a train set of data generated by the RNN on 2400 trials (20 trials for each out of 6 *×* 5 *×* 4 = 120 possible stimulus conditions). For all plots, we first generated new data, changed to an orthogonal basis using **U**^⊤^**M** followed by subtracting with the mean ***µ*** and an orthonormal rotation with **W**_full_.

For analysis of the RNN dynamics (perturbations and fixed points), we transformed the RNN’s latent dynamics such they are expressed in this basis. Take 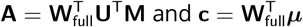. Under the change of coordinates **z**_*new*_ = **Az** − **c**, the RNN parameters are updated as:

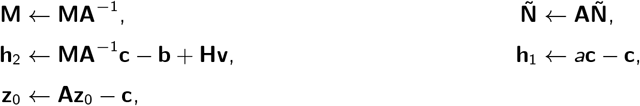

with the RNN now described by:

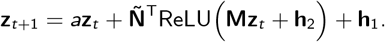

In Fig 2, we plot the variance of the delay activity per component. Because the (fixation) input is constant across conditions during the delay, and because the latent trajectories were expressed in an orthonormal basis, the variance captured in the latent variables **z** corresponds directly to variance in RNN activity **x**.

### onditional fixed points

To find fixed points conditioned on a subset of clamped latent coordinated, we partitioned the latent states into *R*_*c*_ *clamped* and *R*_*m*_ *moving* components:

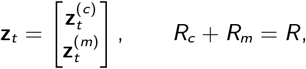

Where 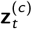 is clamped and 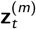 evolves according to the dynamics. We can treat the fixed coordinates as input or bias to the moving latents. Denoting, with …^(*m*)^ and …^(*c*)^ the columns or rows corresponding to moving and frozen coordinates, we obtained a set of *R*_*m*_ equations describing only the moving latents:

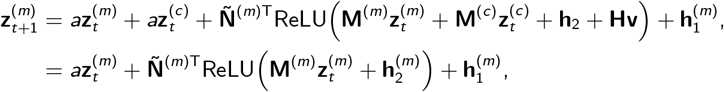

with effective offsets

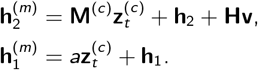

Note we here assumed input **v** to be constant, allowing us to incorporate it as a bias term.For fixed 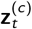, the conditional fixed points are defined by solutions to

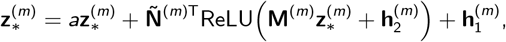

The full latent fixed point is then reconstructed as

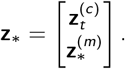

We identified fixed points by simulating the clamped RNN for 7 seconds (140 timesteps of 50 ms) while maintaining the fixation input. The latent state at the final timestep was extracted, perturbed with uniform noise, and used as the initial condition for the heuristic SCyFi algorithm [46].

The SCyFi algorithm is designed to compute fixed points in piecewise-linear systems, such as the ReLU-based RNNs used in this study. In piecewise-linear systems, the latent space is partitioned into regions, each governed by its own linear dynamical system. Given a latent state, SCyFi starts by obtaining the region in which that latent state resides, and computes the fixed point of the corresponding linear system. The resulting candidate fixed point may either lie within the same region, in which case it is a valid fixed point of the system, or fall into a different region. In the latter case, SCyFi reinitializes to the new region and computes the fixed point for the new corresponding linear system. This process is repeated until either a valid fixed point is found (within the current region) or a maximum number of iterations is reached.

### Perturbation analysis

For the perturbation analysis, we simulated the RNNs and applied an impulse current to all RNN units 0.5s. after the last stimulus input. Let 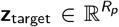 be the target in latent space, where *R*_*p*_ denotes the amount of latents we want to perturb. We can define the perturbation 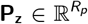 to latents 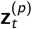, or equivalently, the corresponding perturbation **P**_**x**_ ∈ ℝ^*N*^ to units **x**_*t*_ as:

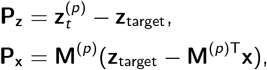

where **M** is assumed to be orthonormal (see above) and **M**^(*p*)^ picks out the columns corresponding to the latents we want to perturb. For Fig. 4d-f perturbation targets were sampled uniformly from a box spanning the delay-period range of each stimulus subspace coordinate, expanded by *×*1.2. Movement was integrated over the next 0.75s to compute the correlation coefficients in Fig. 4e and f.

To model optogenetic-style perturbations (Fig. 5b; right), we constrained perturbations to act with uniform intensity on a subset of recurrent units. We can think of the entries in the dense perturbation vector **P**_**x**_ as computed above, as quantifying the contribution of each neuron to the desired latent-space displacement. We therefore selected the top 5% of units with the largest magnitudes in **P**_**x**_, giving us a sparse mask **m** ∈ *{*0, 1*}*^*N*^. Restricting the perturbation to a binary subset of neurons results in an unscaled latent-space perturbation 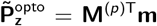 that is generally not perfectly aligned with the target displacement **P**_**z**_. We determined the scaling *c* by minimizing the reconstruction error 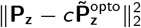 via orthogonal projection. The resulting scaling factor and the corresponding perturbation applied to the recurrent units are given by:

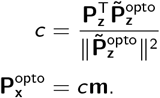

### Behavioral decoding

We trained a shared 6-class logistic regression (with *l* 1 regularization *C* = 1; [76]) to decode the macaques’ responses on (standardized) latents inferred from the macaques spikes using our sequential Monte Carlo framework. We averaged latents over a 0.25 s window ending .05s after the macaques’ responses, and fit the same decoder to the responses corresponding to each sequence position. The decoder trained on inferred latents was then applied to generated latent trajectories, with time windows defined based on the macaques’ average response times.

### Hand-constructed models

To verify our methods, we implemented RNNs that maintain memory of one angular variable *θ* using either attractor or transient dynamics. We used methods from Beiran et al. [41] for constructing low-rank RNNs with Gaussian populations. The models were simulated using:

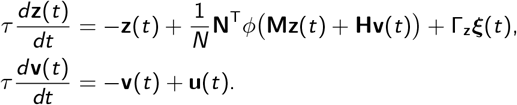

#### Transient dynamics

We constructed a rank-20 recurrent neural network with *N* = 500 units, with *ϕ*(**x**) = ReLU(**x**) and Γ_**x**_ = **M**. The dynamics consisted of two sets of 10 transient modes: one encoding sin(*θ*) and one encoding cos(*θ*).

The connectivity matrix **M** was initialized with i.i.d. Gaussian entries. To generate sequential dynamics, we imposed a feedforward chain within each set of modes. Let **D** ∈ R^*N×*20^ be a random Gaussian matrix. The left connectivity vectors for one set of modes were defined as:

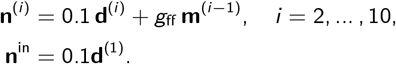

The output mode accumulated contributions from all previous modes:

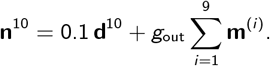

The input weights **I** ∈ R^*N×*2^ were aligned with the first mode of each set:

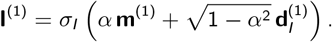

Dynamics were simulated using Euler-Maruyama integration with time constant *τ* = 0.3, time step Δ*t* = 0.05, and Gaussian noise of amplitude *σ* = 0.005 injected in the low-rank subspace.

#### Approximate ring attractor

We constructed a rank-2 RNN with *N* = 600 units and activation *ϕ*(*x*) = ReLU(*x* + 2). Following Beiran et al. [41], the network was partitioned into *P* = 6 equally sized populations, each associated with a preferred angle

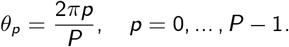

The connectivity vectors were defined using structured population means plus Gaussian noise. For each population

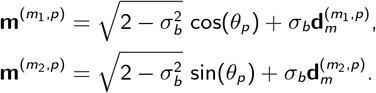

The corresponding left connectivity vectors were

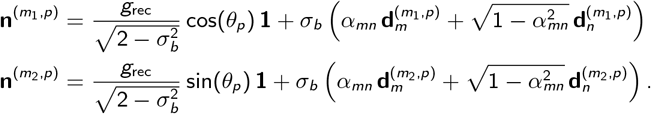

Parameters were set to *σ*_*b*_ = 0.2, *g*_rec_ = 3, and *α*_*mn*_ = −0.5. The positive ReLU bias and recurrent gain push the dynamics away from the origin, while the negative alignment *α*_*mn*_ stabilizes activity along the ring. The input matrix **I** ∈ ℝ^*N×*2^ was aligned with the connectivity modes:

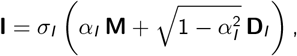

where *α*_*I*_ = 0.75, *σ*_*I*_ = 6, the time constant was *τ* = 0.2, and Gaussian noise with amplitude *σ* = 0.1 was used.

### Dataset for student-teacher setups

To verify that we can in principle recover transient dynamics, we constructed a dataset using the hand-constructed model with transient dynamics. Task input for a total of 6000 trials of length 5s was generated. Each trial had one stimulus presented: **u**_*t*_ = [cos(*θ*), sin(*θ*)], with *θ* drawn uniformly from 6 stimulus conditions. The stimuli were applied for 250 ms, following a random onset of 500 − 700 ms. The RNN activity was mapped to spikes via

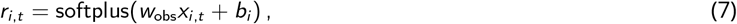

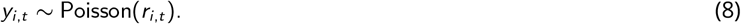

The gain *w*_obs_ and offsets *b* were set so that the resulting spike trains approximately matched the *marginal* statistics of the experimental data (mean firing rates, inter-spike-interval distributions, and pairwise spike-count correlations). In particular we chose, for the transient dynamics: *w*_obs_ = 1.0 and *b*_*i*_ *∼ N* (−0.25, 1.0), and for the attractor dynamics *w*_obs_ = 0.1 and *b*_*i*_ *∼ N* (−0.1, 0.5). Neurons and trials were partitioned into 8 disjoint sessions. Within each session, trials were divided into 75% training and 25% validation sets.

### Session-consistency loss for student-teacher setups

In the student-teacher setups, we found that latent axes were not consistently aligned across sessions in the students with rank 2 or 3: the same stimulus could be represented in different coordinates (e.g. cos *θ* in *z*_1_ for one session and in *z*_2_ for another). This was fixed by the consistency loss described here which encourages a shared latent geometry across sessions. Note that we did not apply this loss for any model fit to macaque data. We hypothesize that the alignment problem is more likely to appear in the student-teacher settings here, where the ground-truth dynamics are more low-dimensional and highly symmetric.

Concretely, during each training batch, we computed for each of the presented stimuli *s*, the mean latent vector 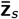 over trials for a fixed delay window of .5s, starting .5s after stimulus offset. We maintained a session-aggregated reference centroid **c**_*s*_, initialized from the first batch in which *s* appeared, then updated with exponential momentum, 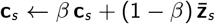, (with *β* set to .9). The auxiliary loss was

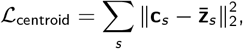

added to the variational objective with weight 50 for the rank-2 RNN fit to the attractor teacher model, and weight 1 for the rank-3 RNN fit to the transient teacher model. The reference centroid was updated after computation of the loss (after which gradients were also detached).

## Supplementary

**Supplementary Figure 1.**
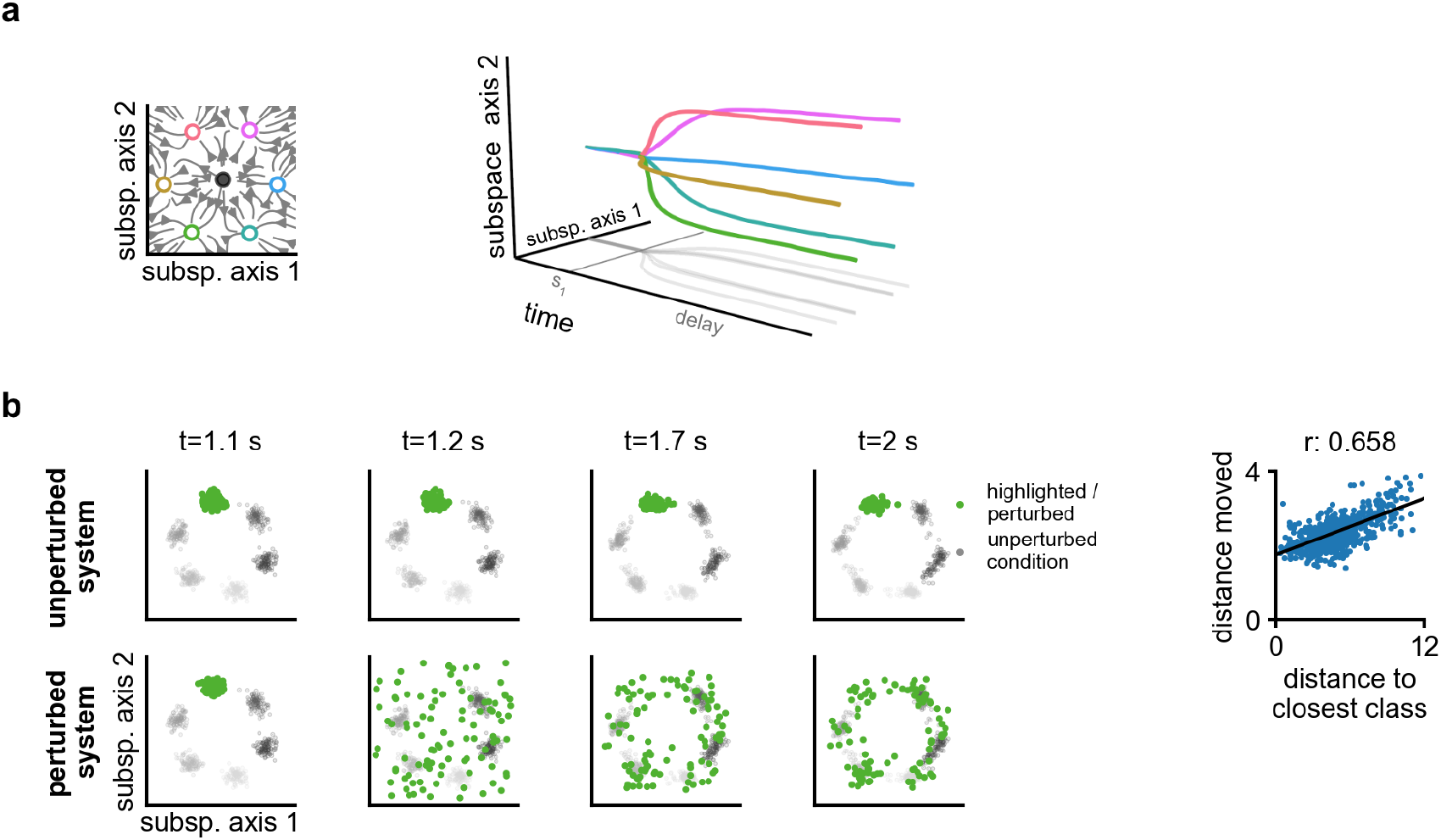
Ring attractor model. **a)** We constructed a rank-2 ring-attractor RNN, with 6 stable fixed points (left). Example trajectories for 6 different stimulus inputs are shown on the right. **b)** Perturbations reveal the underlying attractor structure. We perturb one of the stimulus conditions (green). After perturbation trajectories converge to one of the 6 stable fixed points.

**Supplementary Figure 2.**
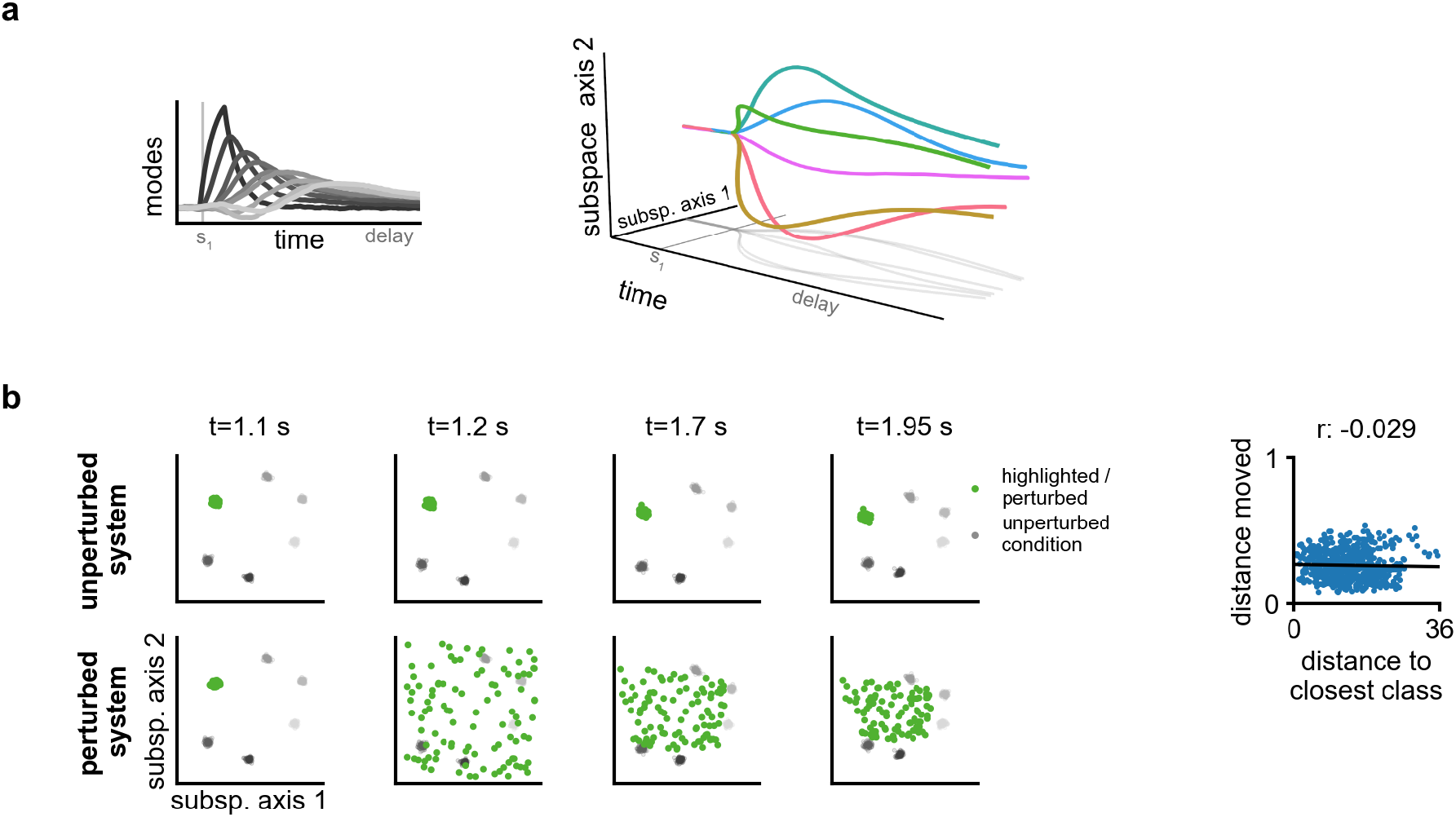
Transient dynamics model. **a)** We constructed a rank-20 RNN, with 2 sets of 10 modes that sequentially activate (left). One set of modes maintains sin of the angular stimuli, and one set maintains cos. Example trajectories for 6 different stimulus inputs are shown on the right, with the subspace axis estimated similar to the main text. **b)** Perturbations again reveal the underlying attractor structure. We perturb one of the stimulus conditions (green). After perturbation trajectories do not move towards the different class means, indicating no evidence for a multi-stable attractor.

**Supplementary Figure 3.**
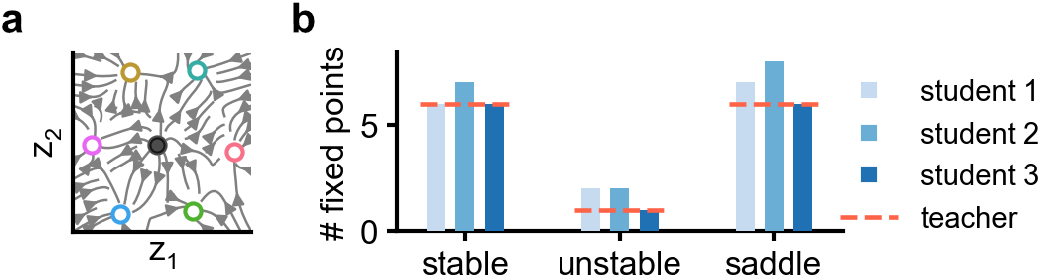
Student-teacher setup with attractor dynamics. **a)** We fit student models with rank 2 to data generated from the teacher model in Fig. S.1. Flow fields from the students match that of the teachers quantitatively. **b)** Generally the students (3 seeds) are able to capture the multi-stable, approximate ring attractor of the teacher, however the students do sometimes find one additional spurious stable fixed point on the attractive manifold.

**Supplementary Figure 4.**
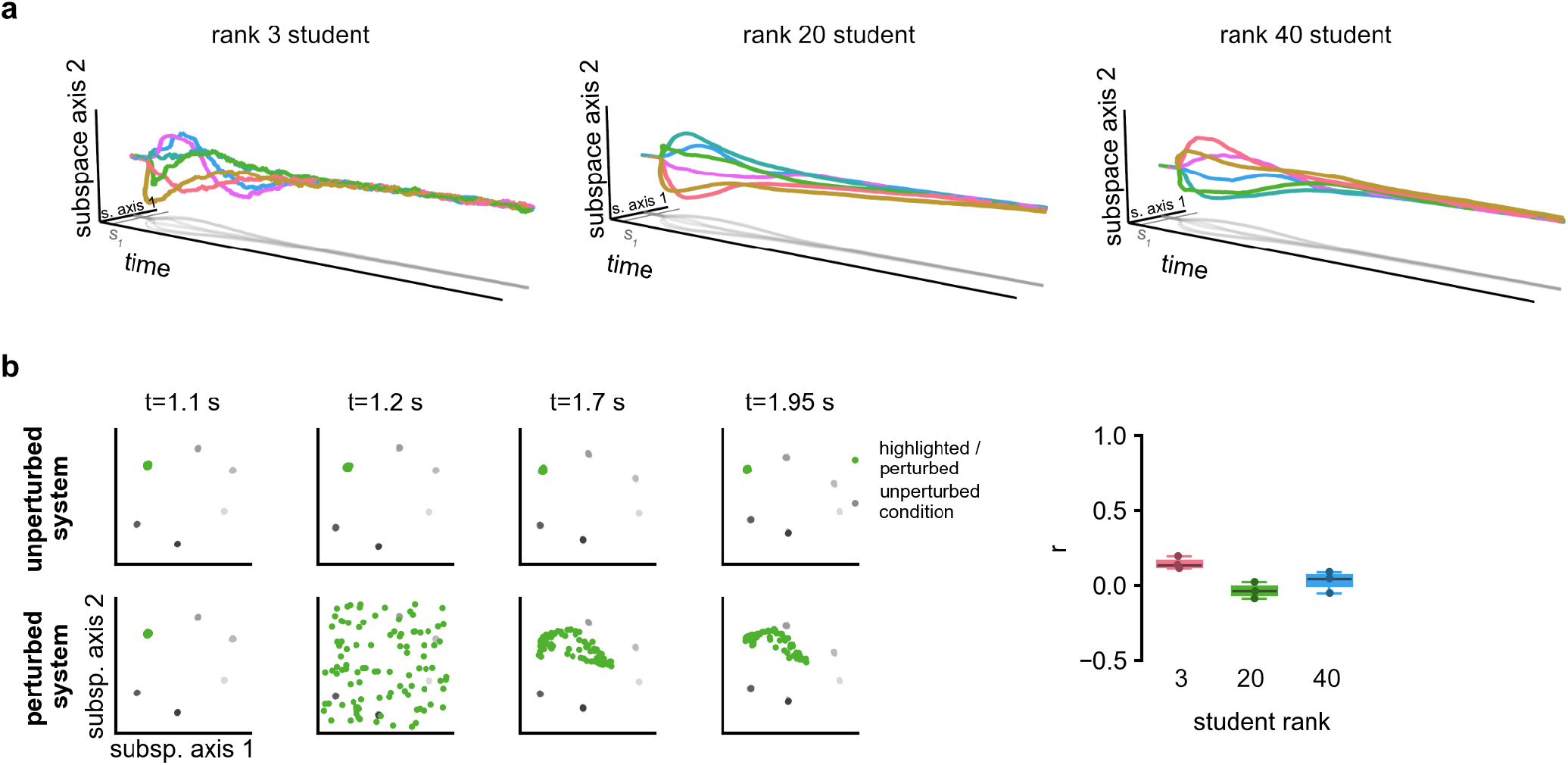
No spurious attractors in student-teacher setup with transient teacher. **a)** We fit student models with rank 3, 20 and 40, to data generated from the rank 20 teacher model in Fig. S.2. In particular the rank 3 student case is interesting as based on Qian et al. [68], we might expect to find spurious attractors when the student does not have the capacity to match the teacher. In order to check whether or not the models learn transient dynamics, we first simulate the RNNs for an extended time after stimulus onset. For all models the latent dynamics collapse, indicating the stimulus maintenance was indeed transient. **b)** We also performed perturbation analysis. We plot one example perturbation on the left, and next, similar to Fig.4, we compute the correlation coefficient over distance moved compared to how far the perturbations pushed the trajectories away from the class means. Across three seeds per student rank, we do not find any strong support for attractor dynamics.

**Supplementary Figure 5.**
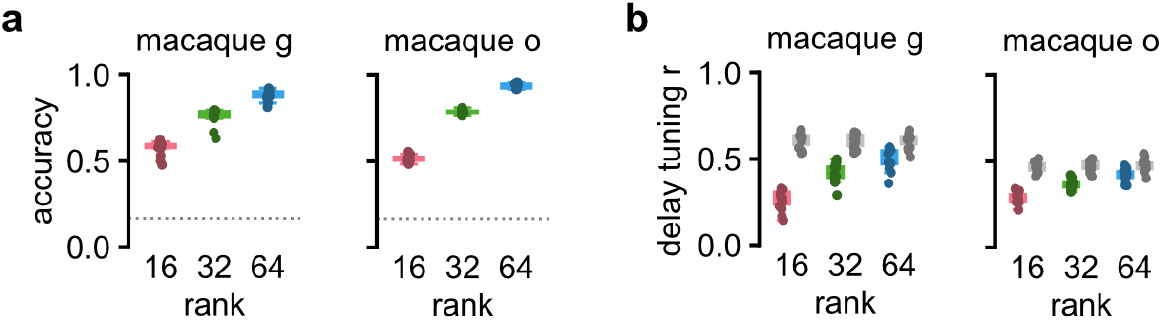
Rank 64 is needed to capture stimulus-conditioned delay activity. Low-rank RNNs were fit at ranks 16, 32 and 64 (5 models per animal; 15 sessions). **a)** Within-session decoding of presented item identity from inferred late-delay latents. Spread is shown over sessions (data was first averaged over models and position). **b)** Stimulus-conditioned delay tuning. For each unit and sequence position, delay rates were first averaged over trials of each item at a particular sequence position, to get a 6-dimensional tuning vector. This was then correlated with the corresponding tuning vector obtained from validation data. Finally all tuning vectors *r* ‘s were averaged over units and positions. Colored boxes indicate model vs validation *r* and gray boxes train vs. validation (note that the gray boxplots are slightly different between ranks as train/validation splits were unique for each model).

